# Different localization of fluorescently labeled N- and C-termini of nucleolin variants in human glioblastoma cell culture

**DOI:** 10.1101/596916

**Authors:** Dmitri Panteleev, Nikolai Pustogarov, Alexander Revishchin, Dzhirgala Shamadykova, Sergey Drozd, Sergey Goryanov, Alexander Potapov, Galina Pavlova

## Abstract

Nucleolus-oriented protein nucleolin plays a significant role in the life of a normal mammalian cell. However, nucleolin is also actively expressed in cells of malignant tumors. At the same time, its expression in different types of cancer is significantly increased compared with normal cells. It is interesting that nucleolin localization often varies in tumor cells, namely in the cytoplasm and on the cell membrane. This fact is considered to be a poor prognostic indicator. This work is devoted to the study of the distribution of nucleolin in human glioblastoma cells. Glioblastoma is one of the most aggressive malignant tumors with an absolutely unfavorable prognosis. These tumors have a high proliferative potential, but in addition they are often characterized by invasive properties. Research on lineage cells does not let to fully study these properties of glioblastoma, since lineage cells are very different from the actual tumor. In our study, we used two primary cell cultures of human glioblastoma with varying degrees of invasiveness of the original tumors. The main interest was directed at studying the localization of nucleolin and its correlation with the invariability of the N- and C-termini of the corresponding protein. Particular attention was paid to the significance of the unaltered C-terminus of nucleolin for its distribution in the cells of transplanted human glioblastoma cultures derived from patient tissues.

The aim of this work is to find the relationship between the deformation of the N- or C-terminal sequences of nucleolin and its localization.We showed that in glioblastoma cells, with a high degree of invasion, nucleolin is found in the cytoplasm and close to the cell membrane, and the distribution of nucleolin with undeformed C and N-terminal does not match.

## 1. Introduction

Nucleolin is a multifunctional protein that has a number of important functions in the vital activities of the cell. For example, its involvement in chromatin remodeling, mRNA stabilization and translational modulation has been shown [Fonseca et al 2015; Fonseca et al 2017]. Nucleolin studies have shown that its N-terminal domain regulates transcription and maturation of ribosomal RNA [Roger et al 2003]. Its central region containing 4 RNA-binding domains has been shown to participate in regulation of the stability of mRNA or preribosomal RNA [Sengupta et al 2004; Serin et al 1996], whereas its C-terminus interacts with ribosomal proteins in regulation of the RNA translation [Abdelmohsen et al 2011; Bouvet et al 1998]. Such diverse and crucial functions of nucleolin indicate the importance of this protein for the functioning of the cell. At the same time, the level of nucleolin has been found to increase in actively proliferating cells (Hsu et al 2015), which is important both for the normal cells of the organism and for pathologically multiplying cells of malignant tumors. In recent years, a large number of articles have been devoted to the study of nucleolin in tumor cells. It has been shown that during the development of various types of malignant tumors nucleolin interacts with EGFR, Ras and Sp1, and this interaction leads to active tumor growth [Hsu et al 2015; Hung et al 2014]. In addition, in some types of malignant tumors, extranuclear cytoplasmic nucleolin has been shown to correlate with poor prognosis. A similar pattern has been found in lung and esophagus cancers [Qi et al 2015; Xu et al 2016]. Nucleolin has also been shown to play an important role in breast tumors (Moura et al 2012) and to be even associated with tumor angiogenesis [Christian et al 2003]. Thus, nucleolin is certainly of interest either as a diagnostic indicator or as a potential therapeutic target of malignant tumors.

When studying the expression of nucleolin in the adult human brain, this protein was shown to be extremely important for the development and existence of nerve cells. NCL has a fundamental role in mediating signals by extracellular matrix molecules and furthermore contributes to the differentiation and maintenance of neural tissue. NCL has been found to be involved in the signal transduction from extracellular matrix molecules and in the differentiation and maintenance of the viability of structured nervous tissue of the brain [Kibbey et al 1995]. When studying the expression of nucleolin in gliomas, it has been shown that the tissue of the most aggressive stage of glioma, glioblastoma derived from patients, over-expresses nucleolin in comparison with the non-pathological samples of the brain (Balça-Silva et al 2018). The increased malignancy of glioblastoma has been shown to correlate with high cell density and with overexpression of nucleolin. The authors have shown that this excessiveness of nucleolin is also associated with a change in the localization of the protein: it is mainly found in the cytoplasm and near the cell membrane [Galzio et al., 2012]. At the same time, blocking nucleolin (for example, with N6L) has been shown to result in a significant decrease in the viability of human glioblastoma cells [Benedetti et al., 2015]. The main conclusion of these studies is the fact that nucleolin, as a diagnostic marker, can be used not only as a factor with increased expression, but also as a factor that changes its localization during malignant processes. Galzio et al. argued that an increase in membrane-localized nucleolin could serve as a poor prognosis indicator and correlate with the level of glioma malignancy [Galzio et al., 2012]. In addition, in the study of cell lines derived from aggressive types of cancer, a large number of shortened splice variants of nucleolin have been identified, which made it possible to suggest a correlation between cell malignancy and increase in diversity of short forms of nucleolin [Hsu et al 2015]. For example, two nucleolin variants of 72 and 55 kDa have been found in lung cancer cell lines, whereas the full-length protein is 110 kDa [Kito et al 2005; Hsu et al 2015]. It should be noted that the authors of these works devote considerable attention to the N-terminal sequence of nucleolin but pay little attention to its C-terminus. In any case, since there is still no clarity on the role and processing of nucleolin, while its importance is obvious, our work is devoted to further research on nucleolin role in human glioma tumor cells. We showed that in glioblastoma cells, with a high degree of invasion, nucleolin is found in the cytoplasm and close to the cell membrane, and the distribution of nucleolin with undeformed C and N-terminal does not match.

## 2. Materials and Methods

### 2.1 Cell Cultures

HEK293 (Human Embryonic Kidney 293), MCF7(Michigan Cancer Foundation-7), MSC (Mesenchymal stem cells of human) and primary glioma cultures were used in this study and grown in Dulbecco’s modified Eagle medium (#C420, PanEco, Russia) or Dulbecco’s modified Eagle medium/F-12 mix (#C470, PanEco, Russia) supplemented with 10% fetal bovine serum (#RYF35911, ThermoFisher, USA). The cells were passaged every week at a split ratio of 1:2. 80% confluent monolayers were transfected using TurboFect transfection kit (#R0531, Thermo Scientific, USA) according to the manufacturer’s protocol. The consequent manipulations were performed 24-48h after transfection.

The research protocol for using samples collected from active duty personnel and civilians who suffered mild TBI in the studies described herein were approved by the Ethics Committee at Burdenko Institute of Neurosurgery (Moscow, Russia). Written informed consent was provided by all research subjects. The privacy of participants was protected using global unique identifiers.

### 2.2 RT-PCR, cloning & sequencing

Total RNA was isolated using TRIzol reagent (#15596026, ThermoFisher, USA) and treated with DNase I (#18047019, ThermoFisher, USA) to remove genomic DNA according to the manufacturer’s protocol. Reverse transcription was performed using a MMLV RT kit (SK021, Evrogen, Russia) and random hexamer primers. The obtained cDNA was used as a template for RT-PCR. PCR was performed with Taq polymerase (PK113S, Evrogen, Russia) or Phusion High-Fidelity DNA polymerase (F530S, ThermoFisher, USA) according to manufacturer’s protocol).

Cloning was performed according to the standard method. For TA-cloning, the pGEM®-T Easy Vector System (#A1360, Invitrogen, USA) was used. Expression constructs were created on the basis of vectors: pTagGFP2-C (#FF191, Evrogen, Russia). Sequencing was performed by the Sanger’s method (Evrogen, Russia). For alignment of sequences, we used available Internet resources (USCS Genome browser, BLAST). For CDS cloning, the following primers were used: F: 5′-AGATCTGACCCCAAGAAAATGGCTCCTCCTCCAAA-3′, R: 5′-GGATCCCTATTCAAACTTCGT-3′.

### 2.3 Rapid Amplification of cDNA Ends

3′ Rapid amplification of cDNA ends was performed using a FirstChoice RLM-RACE kit (Ambion) according to the manufacturer’s instructions. The resulting amplicons were cloned into pGEM-T Easy vector and sequenced. The following primer was used: F1: 5′-AAAGGAAAGAAAGCTGCAA-3′.

### 2.4 Southern blot hybridization of products of 3’RACE

PCR products were subjected to electrophoresis in 1% agarose gel. The agarose gel was incubated in a denaturing buffer (0.5 M NaOH, 2.5 M NaCl) for 5 min. The chromatography paper moistened with the denaturing buffer was covered successively with: two Mini Trans-Blot Filter Papers (#1703932, Bio-Rad, USA), agarose gel, nylon positively charged membrane (#11417240001, Roche, UK), several sheets of filter paper. The transfer was performed overnight. After that, the membrane was neutralized in a neutralizing buffer (1 M Tris, pH 7.4, 1.5 M NaCl). Cross-linking was performed by UV irradiation at a wavelength of 254 nm for 1 min. Hybridization was performed at 63 °C in Church & Hilbert buffer (0.5 M Na-phosphate 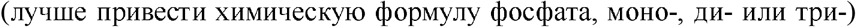, 10 mM EDTA, 7% SDS, pH 7.5). A probe was obtained on the basis of a PCR product using Biotin DecaLabel™ DNA Labeling Kit (#K0651, Fermentas, Lithuania); the probe corresponded to the nucleotides 790-1198 of the NCL transcript (NM_005381.2).

### 2.5 Depletion

Cells of the cell culture transfected with the vGNF vector were lysed in the IP buffer (25 mM Tris, 150 mM NaCl, 1 mM EDTA, 1% NP-40, 5% glycerol, pH 7.4), with the addition of Protease Inhibitor Cocktail (PIK). To remove GFP-carrying proteins, GFP-Trap (#gtak-iST-8, ChromoTek, Germany) was used according to the manufacturer’s protocol. The mixture thus depleted was applied to a column containing M2 anti-flag affinity gel (#F2426, Sigma, USA). The subsequent operations were performed according to the manufacturer’s protocol, with the exception that the elution was performed using X3 Flag peptide (#F4799-4MG, Sigma, USA). Then 5x Non-Reducing Lane Marker Sample Buffer (#1859594, ThermoScientific, USA) was added and Western blotting was conducted.

### 2.6 Western Blot

Western blot analysis was performed by a standard method (Sambrook and Russell, 2001) using a Hybond ECL nitrocellulose membrane (Amersham, USA). PageRuler Plus Prestained Protein Ladder (#SM1811, ThermoScientific, USA) was used as a molecular weight marker. Primary antibodies were used: anti-GFP in a titer of 1:8000 (AB011, Evrogen, Russia), anti-NCL in a titer of 1:7000 (#a300-711A, Bethyl, USA), anti-FLAG M2 in a titer of 1:8000 (#F3165-.2MG, Sigma, USA).

### 2.7 2D Western Blot

The cells were detached from the culture flask with trypsin-EDTA solution (#P039, PanEco, Russia) and washed with PBS. The cells were lysed in O’Farrel lysis buffer supplemented with 1% PMSF and 1x Halt protease inhibitor single-use cocktail (#78430, Thermo Scientific, USA) for 15 min at 4 °C. Then the samples were centrifuged at 15000 g for 10 min and the supernatant was collected. Electrofocusing was performed in glass capillaries with polyacrylamide gel using a mixture of ampholytes 3/10 and 5/8 (#163-1112, #163-1192, Bio-Rad, USA). The second direction is standard SDS 5-10% PAGE followed by staining with Coomassie Brilliant Blue G-250 (#31-4-58-1, Helicon, Russia) or ELISA using anti-NCL antibodies (#a300-711A, Bethyl, USA).

### 2.8 Immunofluorescent staining of cells

Cell staining was performed according to a standard procedure. Fixation was performed with 4% paraformaldehyde for 15 min. As primary antibodies, anti-FLAG M2 were used in a titer of 1:350 (#F3165-.2MG, Sigma, USA). The obtained preparations were analyzed using a Leica TCS SP2 confocal microscope.

Lamin detection was performed using #ab 16048 (abcam, USA) antibodies in 1/400 titer/ The obtained preparations were analyzed using fluorescent Leika20a.

## 3. Results

We selected several primary cultures derived from human gliomas GrIV. Tumors were removed at the Burdenko Institute, Moscow, Russia. Glioblastoma cell cultures were cultivated for 5-15 passages. For comparison, the HEK293 line was taken in the first experiment; this culture is often used as a conditionally normal one. On this selection of cultures, a variety of nucleolin splice variants was investigated by the Southern blot hybridization of 3’RACE products using primer F2. The result (Fig. 1) showed the presence of multiple bands, which indicates the abundance of splice variants of nucleolin in these cultures, whereas the combination of splice variants is more abundant in cancer cells.

**Fig. 1.**
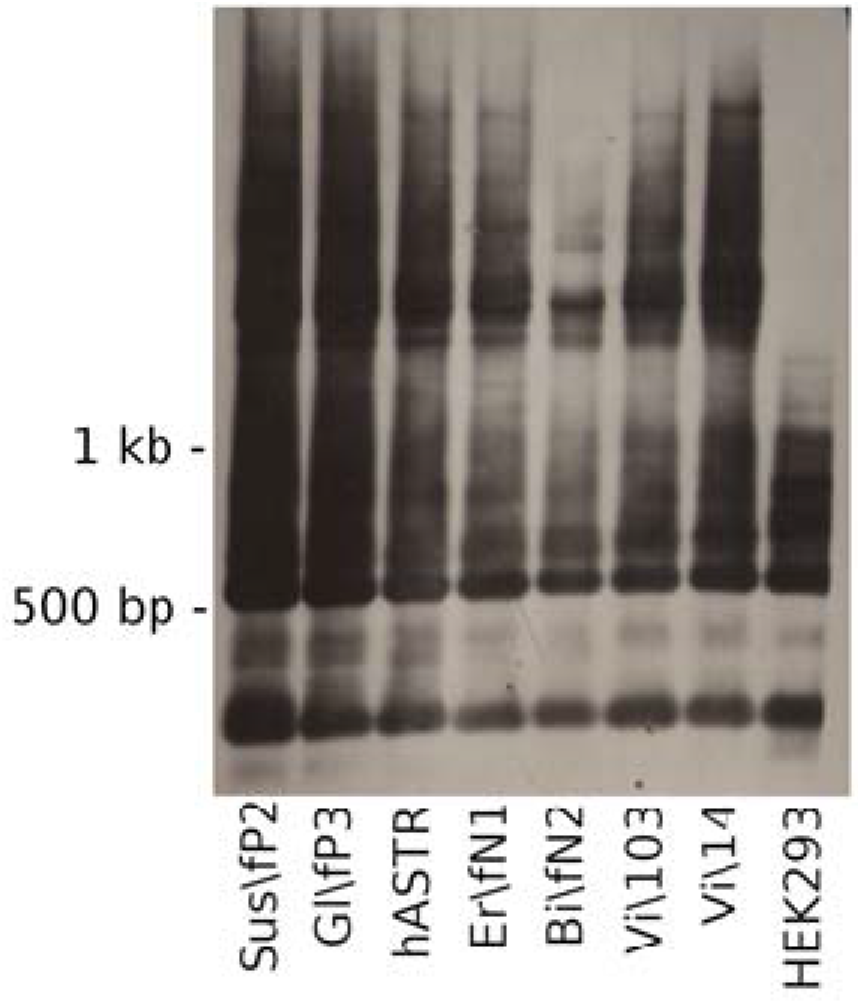
Southern blot hybridization of the products of 3’RACE with cell culture lysates for nucleolin using primer F2. The presence of multiple bands of different mobility is noticeable.

Such a significant increase in alternative splicing may indicate disruptions that result either in the synthesis of a large number of splice variants without an increase in protein variants, or in the expansion of the spectrum of nucleolin proteins.

In order to understand whether nucleolin diversity increases in tumor cells, we investigated the lysates of two cell lines, HEK293 and MCF7, by 2D Western blot hybridization method. HEK293 is considered conditionally normal in all studies. The HEK293 line is derived from human embryonic kidney cells, while MCF7 (Michigan Cancer Foundation-7) is an epithelial-like cell line derived from invasive adenocarcinoma of the human mammary gland ducts. The latter is one of the most common cell lines for *in vitro* research on the molecular biology of cancer. A comparative analysis of these two lines (Fig. 2) showed that not only the diversity of splice variants of nucleolin RNA in malignant tumors increases, but also the diversity of nucleolin proteins.

**Fig. 2.**
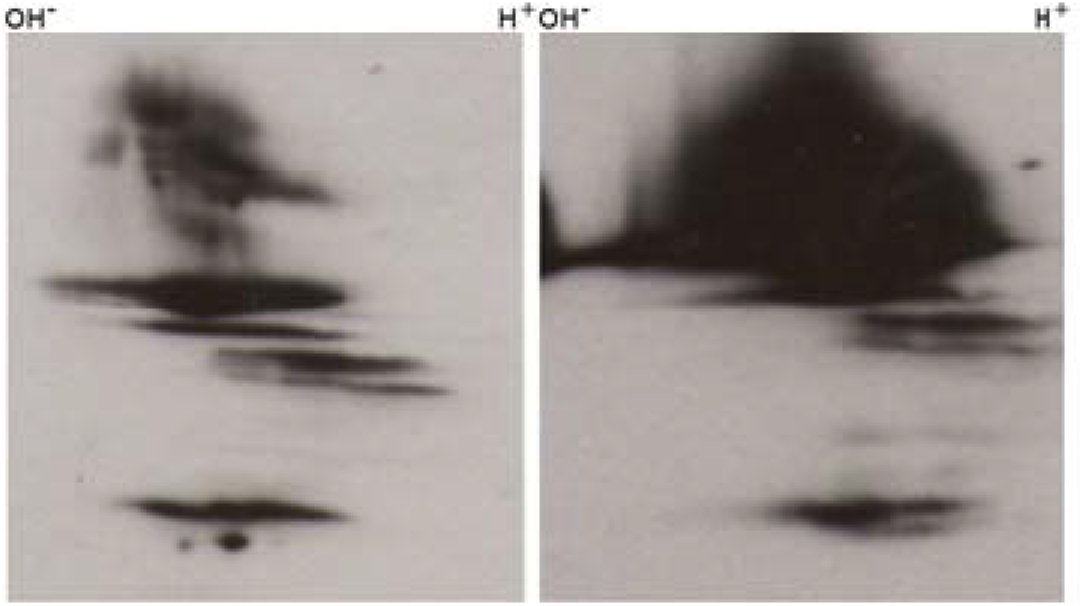
2D Western blot hybridization of the lysates of HEK293 (left) and MCF7 (right) cell lines for nucleolin. The increase in nucleolin variants in the cancer line is noticeable.

Based on data from literature and our preliminary research, we suggested that the presence of an unchanged C-terminus of nucleolin is as important for the correct nuclear localization of nucleolin observed in non-tumor cells, as its N-terminus, which was described in many publications (Xiao et al 2014). Thus, the sequence of the C-terminus can be expected to be disrupted or truncated in tumor cells. Since the field of our research is mainly related to the nervous system, special attention was paid to the study of nucleolin variants in glioma cells. In addition, we chose to use partially transplanted cell cultures derived from post-surgical human glioma tissue, instead of glioma cell lines. Lineage cells are quite different from the cells of the original tumor due to extremely long passaging and immortalization. In our work, two cell cultures, Sh\fP1 and Sus\fP2, were investigated. Sh\fP1 was derived from glioblastoma tumor tissue (GrIV). The tumor was removed from the right motor area of the brain of a 54-year-old man. The duration of the life of the patient after surgery was 1 year and 2 months.

Sus\fP2 was also derived from the glioblastoma tumor tissue (GrIV). The tumor was removed from the left temporal lobe of the brain of a 60-year-old woman. This tumor is distinguished by the fact that the patient in the next 1 year and 8 months had two more surgeries and spreading of the tumor to the deep structures of the brain was found. The duration of the life of the patient after surgery was 1 year and 8 months.

These cultures were analyzed by immunohistochemistry for the proliferation marker Ki67 (Fig. 3).

**Fig. 3.**
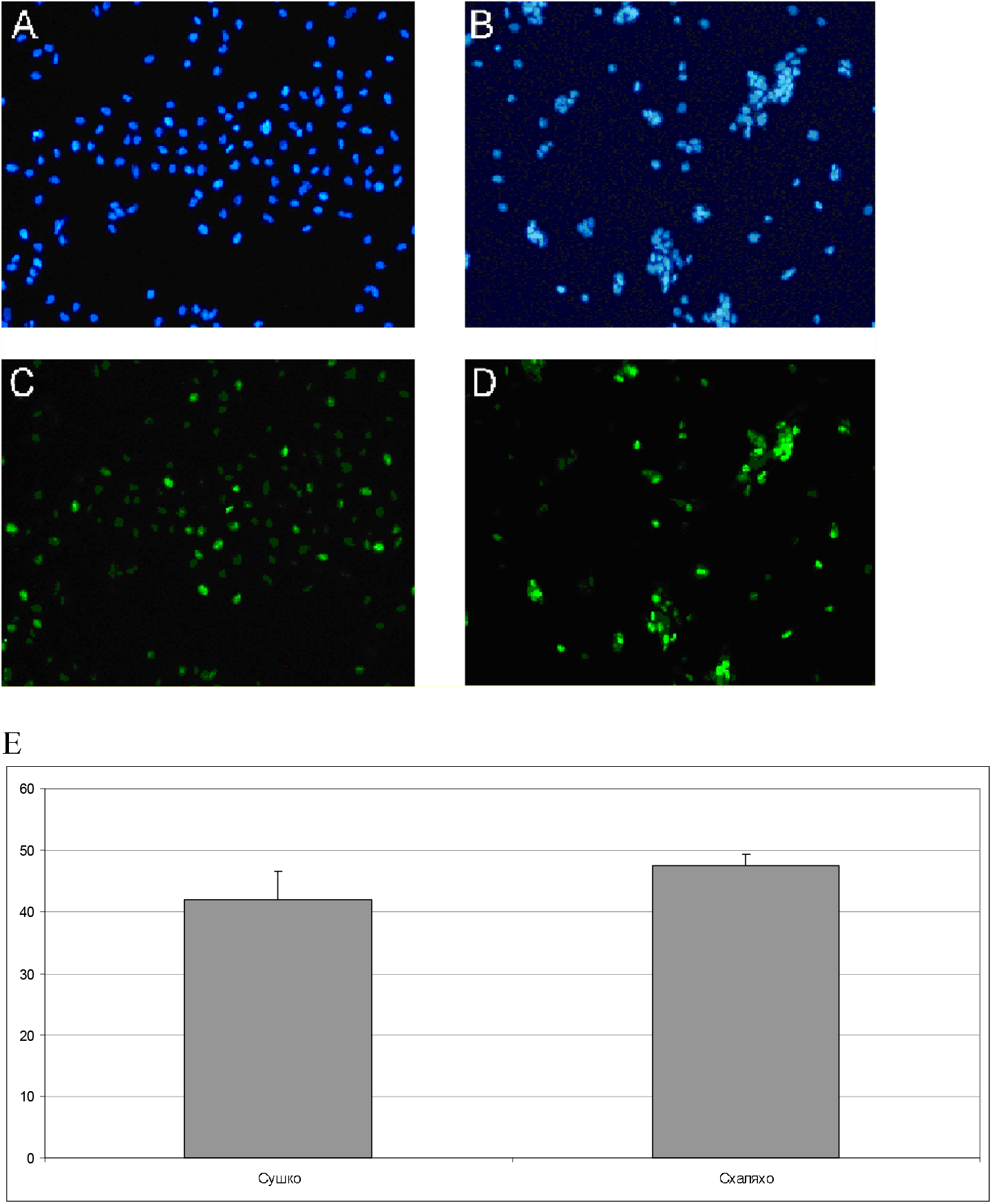
The proliferative activity of Sh\fP1 and Sus\fP2 cultures detected by immunohistochemical staining with antibodies against Ki67. A-D – microphotographs of cell cultures stained with antibodies against Ki67. E – the percentage of Ki67-positive cells in Sh\fP1 and Sus\fP2 cell cultures.

The Sh\fP1 and Sus\fP2 cultures have a high percentage of actively proliferating cells. Thus, we can assert that we have two cultures derived from glioblastoma tissues and having a similar high proliferative potential.

The Sh\fP1 and Sus\fP2 cultures were also analyzed by RT-PCR for the expression of a number of markers (Fig. 4). For comparison, the MCF7 line was used, the expression was normalized against MSC adopted as the norm. The Sus\fP2 culture was found to have an elevated expression of MAP2, PDGFA1, PDGFA2, PDGFB, and also to a lesser extent CDK6, GFAP, MELK and OLIG2.

**Fig. 4.**
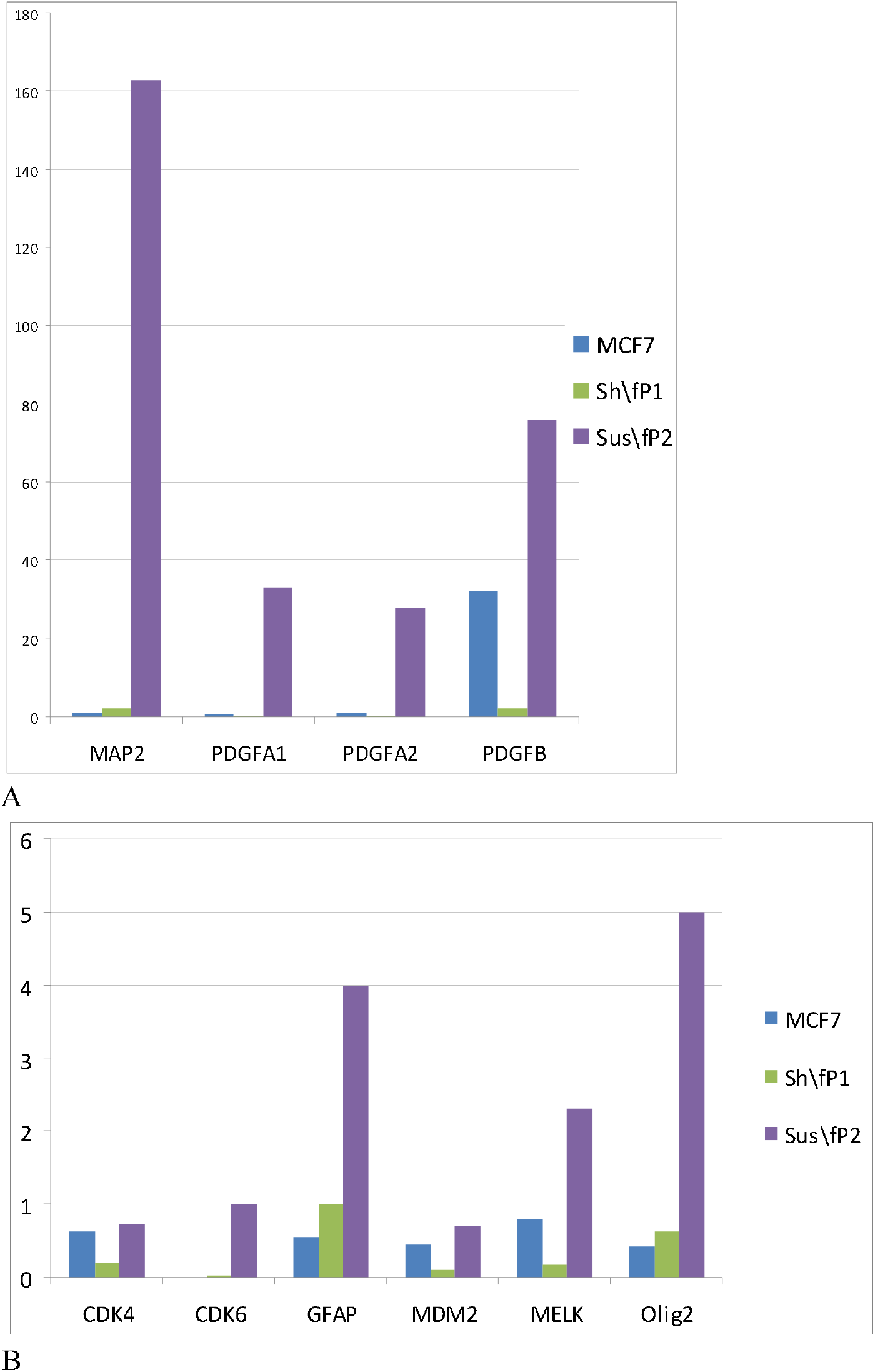
Normalized expression of target genes in human glioblastoma cell cultures (Sh\fP1 and Sus\fP2) and MCF7 line of human breast adenocarcinoma. Expression is normalized against MSC.

It is interesting that the MAP2 marker is differently expressed in the cells of two aggressive forms of glioma. A number of researchers found a similar increase in expression in gliomas and suggested that it can be a marker of glioma (Blümcke et al., 2001). Some researchers argue that MAP2 plays a dual role in the progression of cancer, which was studied in non-glioma tumors (Song et al., 2016). However, some authors assert that MAP2 may even be a positive marker of glioma, as shown in (Yi et al., 2018). However, in our work, we do not see a positive prognostic value of increased MAP2 expression for a patient. On the contrary, an increased expression of MAP2 coincides with an increased expression of PDGFA1, PDGFA2, PDGFB, CDK6, GFAP, MELK, which are considered markers of tumor aggressiveness. We would rather agree that the expression of MAP2 is observed in immature glial progenitors as well as in cells of the aggressive form of glioma, and it is likely to be a negative prognosis indicator.

Also of interest is an increased expression of OLIG2 in Sus\fP2 glioblastoma cell culture that has a high degree of invasiveness. A number of studies indicated that glioma cell populations with a high degree of invasiveness demonstrate an increased expression of OLIG2 with a simultaneous increase in expression of the proliferative marker Ki67 and stem cell marker CD133 (Lu et al., 2016; Ligon et al., 2007; Singh et al., 2016).

Thus, an increased expression of these factors may be associated with the aggressive (malignant) behavior of these cells migrating into the brain parenchyma, which leads to an intensive development of relapses metastases.

For further research, we used as a control the MSC culture which is not a tumor cell culture. In a number of studies, the HEK293 line was also used as a potential control. For comparison with another type of tumor, we used the MCF7 line. At first, we repeated the Southern blot hybridization of the 3’RACE products and showed the presence of multiple nucleolin bands of varying mobility (Fig. 5).

**Fig. 5.**
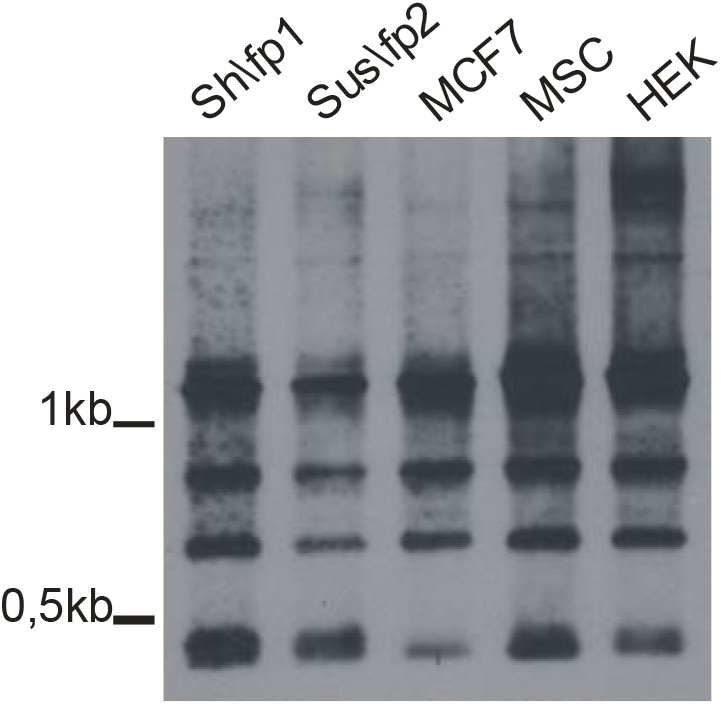
Southern blot hybridization of 3’RACE products from cell lysate for nucleolin using primer F2. The presence of multiple nucleolin bands of varying mobility is noticeable.

Next, we cloned a vGNF construct based on the pTagGFP2-C vector, where the nucleolin gene was tagged at the 5 ‘end with the GFP gene, and at 3’ end with FLAG (Fig. 6). Such a construction may allow us to analyze the distribution of nucleolin with full-length C- and N-termini in the cell and compare their distribution among themselves. This construct was transfected into cells of Sh\fP1, Sus\fP2 and MCF7 cultures. As a control, the MSC culture was also transfected with this vector.

**Fig. 6.**
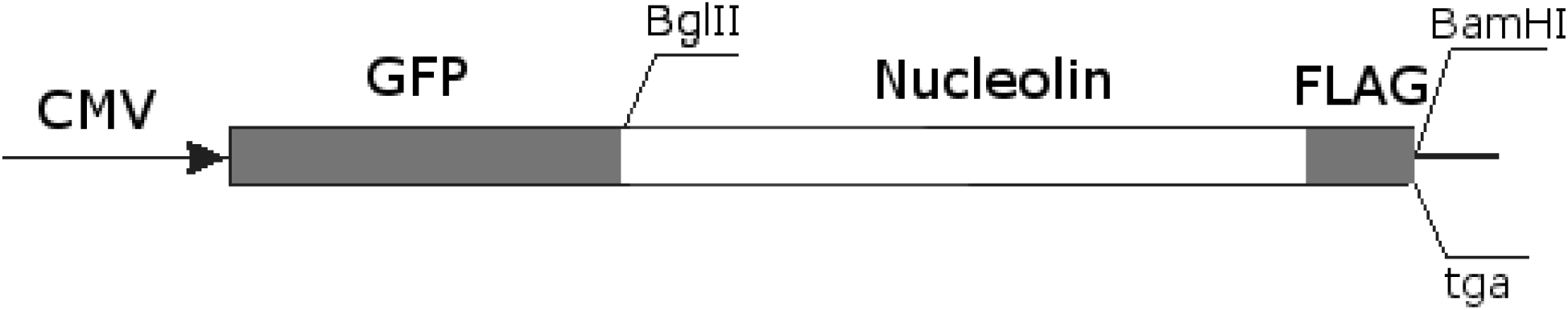
Scheme of vGNF construct used for transfection of cells with subsequent immunocytochemical studies.

As a result, a number of changes were found in the distribution of nucleolin in tumor cells as compared with normal cells (Fig. 7). There is also a difference in the distribution of N-terminal and C-terminal nucleolin in the cells. For instance, in cells of the human Sh\fP1 glioblastoma culture with a low degree of expression of the above markers that correlate with tumor aggressiveness, nucleolin was localized in the nucleus and nucleoli, moreover, the distribution of N-terminal and C-terminal labeled nucleolin coincided. The same pattern was observed in the MSC cells, where the distribution of N-terminal and C-terminal labeled nucleolin was also observed in the nucleus and nucleoli. However, when comparing these two figures, the difference between their distributions is noticeable. The cells of the MSC culture have a clear oval nucleus and the nucleolin is predominantly located in the nucleoli and nuclear envelope, while its distribution in the body of the nucleus itself is insignificant. In the Sh\fP1 glioblastoma cells with a lower expression of tumor markers, and, according to the patient’s case history, a lower invasiveness, the predominant distribution of nucleolin is observed in the body of the nucleus. Thus, there are differences in the nucleolin distribution between normal cells and glioblastoma cells with low invasiveness. But the nucleolin distribution in the cells of the aggressive actively proliferating human glioblastoma cell culture Sus\fP2 (the original tumor, from which the Sus\fP2 culture was obtained, had a high degree of invasiveness) is much more different. In the Sus\fP2 cell culture, we found that nucleolin is predominantly localized on the periphery of the cytoplasm of cells, concentrating at its outer border. It should also be noted that the cell nuclei are either completely devoid of nucleolin or contain only small fragments of it. In any case, tumor cells differ from normal or conditionally normal cells in dislocation of nucleolin in the nucleus and its predominant localization in the nucleolus. If we analyze where the green and red fluorescence signals coincide, that is, the distribution of nucleolin with both N- and C-termini, then we can confidently say that in normal and conditionally normal cells a complete coincidence of green and red luminescence signals is observed. Such a coincidence suggests that normal and expected version of nucleolin is synthesized. An analysis of tumor cultures demonstrates a significant discrepancy in the localization of red and green fluorescence signals. For example, a significant discrepancy is found when analyzing Sus\fP2 and MCF7 cultures. Nucleolin with an undeformed N-terminus predominantly loses its association with the nucleolus and is localized in the nucleus, whereas in the Sus\fP2 culture it is found even in the cytoplasm of cells. Although nucleolin with a non-deformed C-terminus often retains localization in the nucleolus even in a number of malignant cell lines (MCF7 and Sh\fP1), but it can also lose such localization (Sus\fP2).

**Fig. 7.**
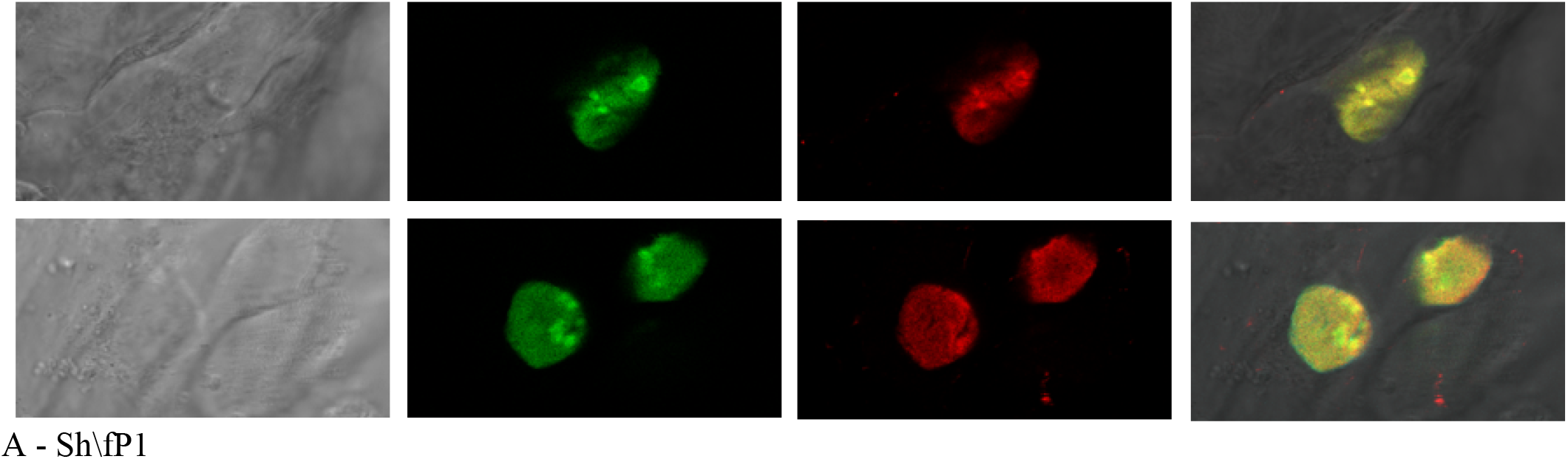
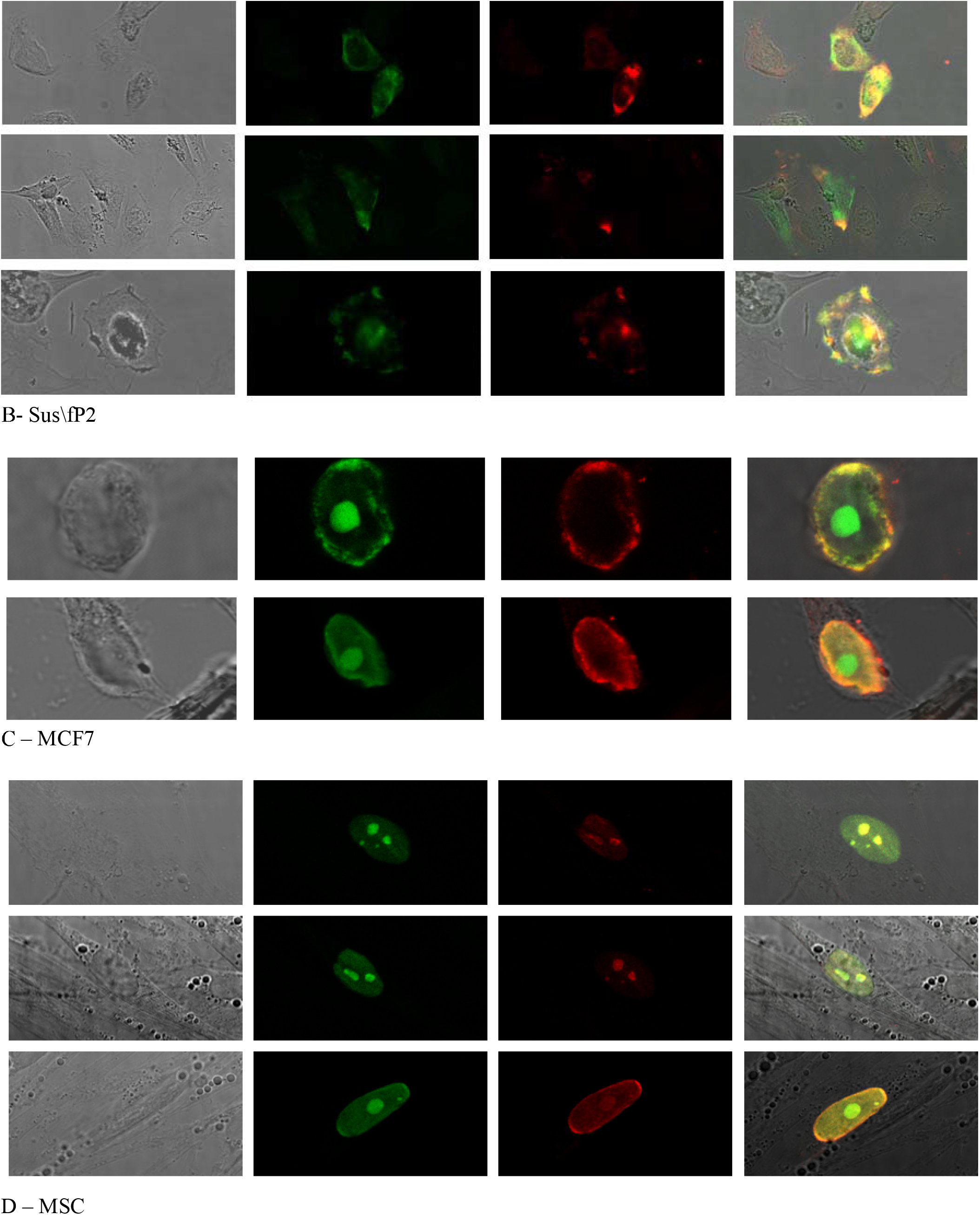
Confocal microscopy of cells transfected with vGNF vector (GFP_NCL_FLAG) followed by staining with antibodies against FLAG (#F3165-2MG, Sigma, USA). The distribution of N-terminal (GFP) and C-terminal (FLAG) tags varied in different cell cultures. A – Sh/fP1 – glioblastoma cell culture Grade IV, B – Sus\fP2 – glioblastoma cell culture Grade IV that has an increased invasive ability, C – MCF7 – human mammary duct adenocarcinoma cell line, D – MSC – human mesenchymal stem cells (control). Scalebar = A, B – 20 μm, C, D – 10 μm.

Lysates of obtained transgenic cultures transfected with the vGNF vector were analyzed by Western blot analysis using anti-GFP (A – N-terminus of fusion protein) and anti-FLAG antibodies (B – C-terminus of fusion protein) (Fig. 8). A number of samples demonstrated a multiplicity of bands. It is worth noting that this multiplicity is found in Sus\fP2 and Sh\fP1 human glioblastoma cell cultures when anti-GFP antibodies are used, whereas this picture is different in mobility and in the presence of additional bands when anti-FLAG antibodies are used. The largest diversity of bands was observed in the sample of the Sus\fP2 cell culture that had an increased capability of invasion. Interestingly, there is a discrepancy between the fluorescence signals in the cells of these cultures as detected by confocal microscopy. These facts indicate the presence of the processes of proteolytic cleavage of nucleolin in the cells with the formation of products that carry only the N-terminal tag and have a characteristic cellular localization specific for tumor cells.

**Fig. 8.**
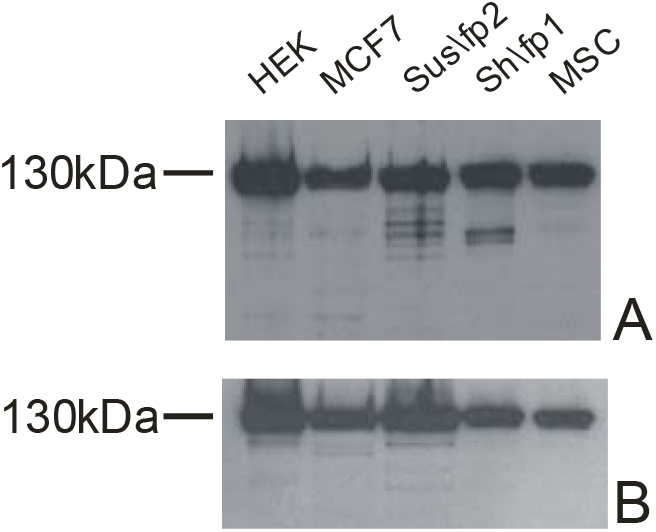
Western blot analysis of lysates of cell cultures transfected with vGNF vector using anti-GFP (A – N-terminus of fusion protein) and anti-FLAG antibodies (B – C-terminus of fusion protein).

To understand how pronounced are nucleolin variants lacking the N-terminus that appear in tumor cells, the lysate of the MCF7 tumor cells was obtained, and then N-terminal depletion was performed. To remove GFP-carrying proteins with the full-length N-terminus, we used the GFP-Trap (#gtak-iST-8, ChromoTek, Germany). With the obtained cell lysate and the depleted lysate lacking proteins with the N-terminus, we performed Western blot hybridization that demonstrated the appearance of a significant number of nucleolin variants that had the C-terminus but were lacking the N-terminus (Fig. 9).

**Fig. 9.**
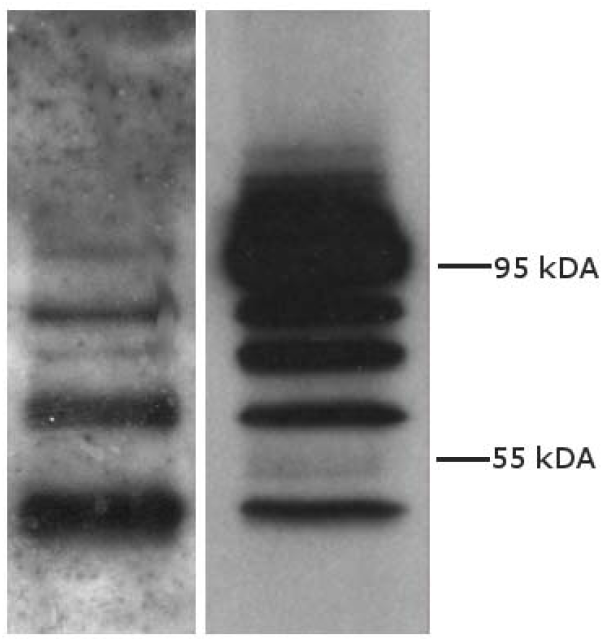
Western blot hybridization of lysates of MCF7 cells (on the right) and proteins remaining after the N-terminal depletion. Several forms of NCL are noticeable. Moreover, the differences in the patterns and ratios of band intensities indicate the presence of forms that do not contain the N-terminus.

Further, we used the 2D Western blot hybridization method to investigate the differences in diversity and concentration of nucleolin with the undeformed N-terminus and nucleolin with the undeformed C-terminus in the actively proliferating glioblastoma culture Sus\fP2 (Fig. 10).

**Fig. 10.**
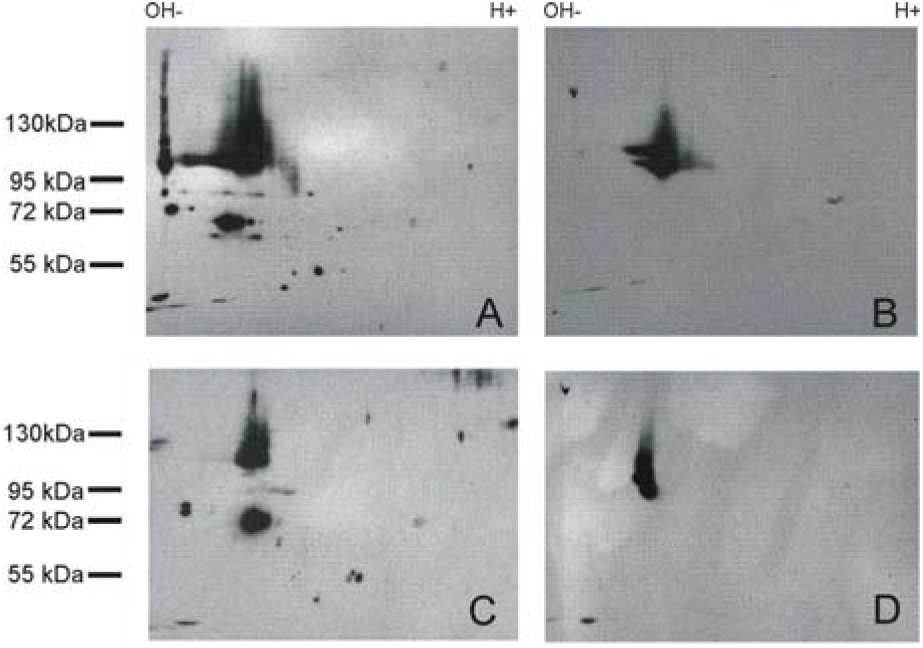
2D Western blot hybridization of the Sus\fP2 (A,B) and MSC (C,D) cell culture lysate. A – antibodies against the N-terminus of nucleolin (#N2662, Sigma, USA). B – antibodies against the C-terminus of nucleolin (#A300-711A, Bethyl, USA). The presence of multiple protein products recognized by antibodies against the N-terminus but not antibodies against the C-terminus is noticeable.

Assessing the localization of nucleolin in cells, we have also used color to lamin, in order to ensure that the localization in a particular case is found in the nucleus or out of it. Since often in tumor cells, especially in Sh \ fP1 and MCF7 cells, large nuclei were found that filled almost the entire cell, therefore such evidence was necessary. It is precisely on the basis of these studies that we were able to convincingly say that in the Sh \ fP1 cells the nucleolin is still localized in the nucleus (Fig. 11)

**Fig. 11.**
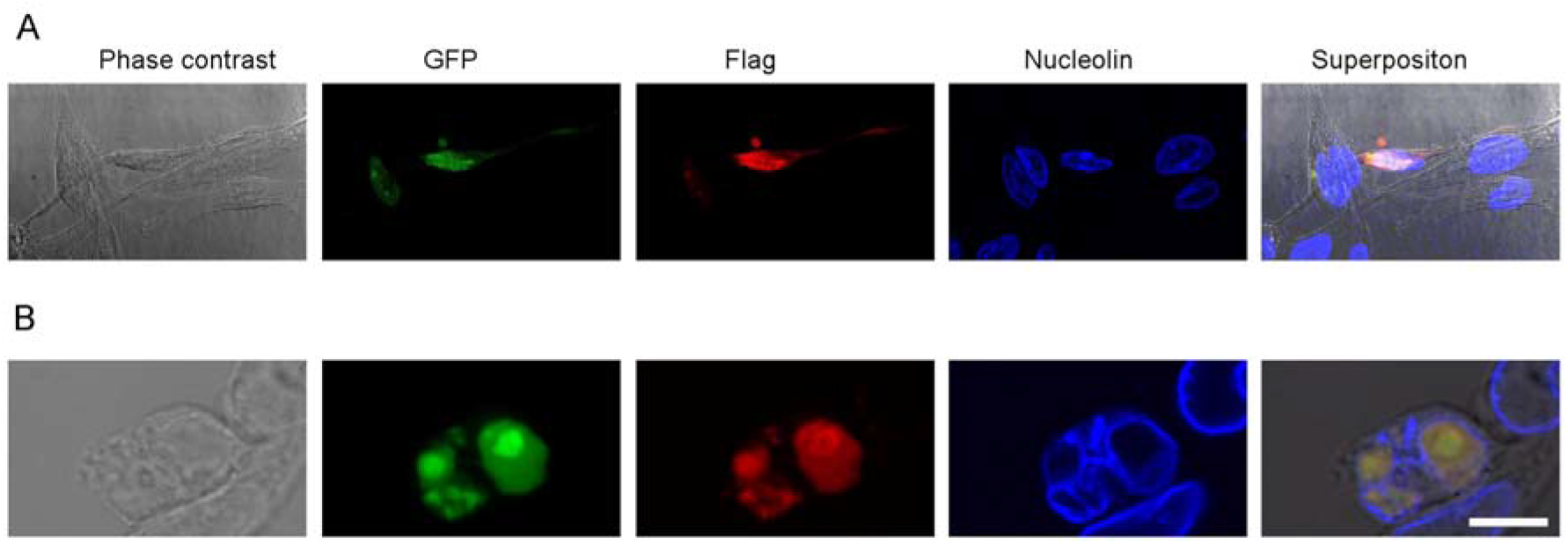
Confocal microscopy of cells transfected with vGNF vector (GFP_NCL_FLAG) followed by staining with antibodies against FLAG (#F3165-2MG, Sigma, USA). A – HEK293, B – Sh/fP1 – glioblastoma cell culture Grade IV.

## 4. Discussion

This study showed a significant predominance of nucleolin variants with the undeformed N-terminus in the actively proliferating human glioblastoma culture Sus\fP2, moreover, the tumor from which the culture was derived had a high degree of migration. This significant predominance of nucleolin with the undeformed N-terminus was manifested both as an increase in the concentrations of identical variants of nucleolin and as an increase in the diversity of nucleolin forms. This result is consistent with the results of immunohistochemistry that showed a sharp increase in nucleolin with localization in the cytoplasm. From these two studies, it can be concluded that nucleolin with the undeformed N-terminus can lose nucleolar localization and move into the cytoplasm. It should be noted that nucleolins with the undeformed N-terminus, in addition to the diversity of forms, demonstrated a variety of localizations. As can be seen in Fig. 7, both nucleolin variants with the undeformed N-terminus (green) localized in the nucleus and nucleolin variants with the undeformed N-termini (green) localized on the periphery of the cytoplasm and along the membrane border are detected. At the same time, nucleolin with the undeformed C-terminus was very rarely found in the nucleus in the Sus\fP2 culture, and if it was detected, it was localized only in the nucleolus, and the majority of nucleolin was observed in the cytoplasm; moreover, the localization of nucleolin with the undeformed N-terminus (green) and with the undeformed C-terminus (red) often did not coincide.

The interest in nucleolin was triggered by a number of studies that showed that this protein is primarily localized in the nucleolar region and is associated with the pre-rRNA (Ginisty et al 1999). Nucleolin has been shown to be involved in ribosome biogenesis, from the structure of rDNA chromatin and regulation of the transcription of RNA polymerase I to the maturation of pre-rRNA and the assembly of preribosomal particles (Ginisty et al 1999). At the same time, a number of studies prove that the ability of nucleolin to interact with RNA and DNA is associated with the post-translational modifications of its N-terminal domain (Cong et al, 2011). Based on such studies, it was suggested that the undeformed N-terminal domain of nucleolin plays an important role in the localization of this protein in the nucleolus (). Our results, without refuting this suggestion, let us argue that the C-terminus is equally important for the normal localization of nucleolin. We observed an increase in the level of nucleolin with the undeformed N-terminal domain in the Sus\fP2 cell culture. Thus, the loss of the normal C-terminal domain (fig. 12) is characteristic of an aggressive glioblastoma culture that is derived from a highly invasive original tumor and has an elevated nucleolin localization in the cytoplasm of the cell.

**Fig. 12.**
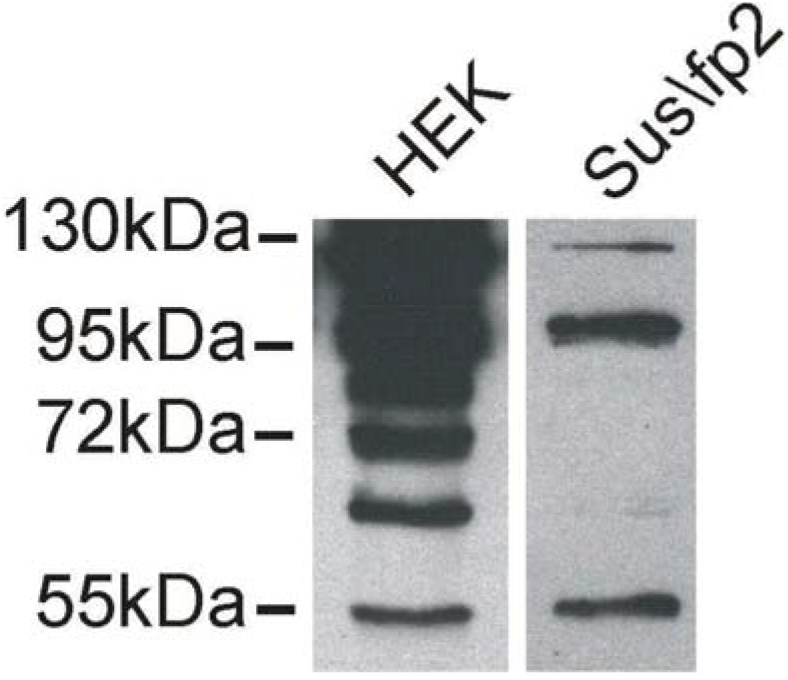
Western blot hybridization of lysates of Sus/fP2 (on the right) and HEK 293 (on the left) proteins remaining after C-terminal enrichment followed by the N-terminal depletion. Several forms of NCL are noticeable. Moreover, the differences in the patterns and ratios of band intensities indicate could correlate with different cell localization (see fig. 7).Stainig with antibodies against FLAG (#F3165-2MG, Sigma, USA).

According to the literature, nucleolin is localized in the cytoplasm; it is assumed to play a significant role in the endocytosis pathway (Legrand et al 2004; Hovanessian et al 2000). At the same time, Losfeld et al. showed that for nucleolin located on the cell surface, an essential requirement is glycosylation of the N-terminus of the protein, that is, the role of the intact structure of the N-terminus of nucleolin is important for nucleolar protein localization (Losfeld et al, 2009). However, we observed in the Sus\fP2 culture both nucleolin with an intact C-terminus on the cell surface and nucleolin with an intact N-terminus that is also localized on the cell surface or in the cytoplasm. Moreover, the localizations of these nucleolin variants often do not coincide, which indicates an increase in the spectrum of shortened nucleolins in the Sus\fP2 cells and the importance of the C-terminus for nucleolar protein localization as well.

In general, a large number of studies have shown that the localization of nucleolin in the cytoplasm of the cell and on the cell membrane is not typical for “healthy” cells. For instance, an increased cytoplasmic expression of nucleolin has been shown to be associated with a worse prognosis for patients with gastric cancer (Qiu et al, 2013), ovarian and breast cancer (Woo et al, 2017), non-small cell lung cancer (NSCLC) (Xu et al, 2017). It is important that the significant role of nucleolin in the control of proliferative properties of cells (Ugrinova et al, 2007) as well as its anti-apoptotic properties that enable tumor cells to successfully multiply and increase the size of a malignant formation (Chen et al, 2012) have been proven. Special attention should also be paid to the work (Garcia et al, 2011), the authors of which suggested that cytoplasmic nucleolin is involved in the metastasis of tumors, which is consistent with our results with the Sus\fP2 culture derived from glioblastoma with a high degree of invasiveness. We cannot claim that this correlation was absolutely proven; this requires more extensive research. But we can argue that the Sus\fP2 human glioblastoma culture derived from tumor tissue with a high degree of invasiveness tends to have a very different distribution of nucleolin in the cells that is characterized by an increase in nucleolin level in the cytoplasm. It should also be noted that while nucleolin level is increased in the cytoplasm, the amount of this protein with the undeformed N-terminus in the cell is much higher than that with the undeformed C-terminus. This study makes us turn our attention to the significance of the C-terminal sequence of nucleolin for its distribution in the cell and its effect on the invasive properties of glioblastoma cells.

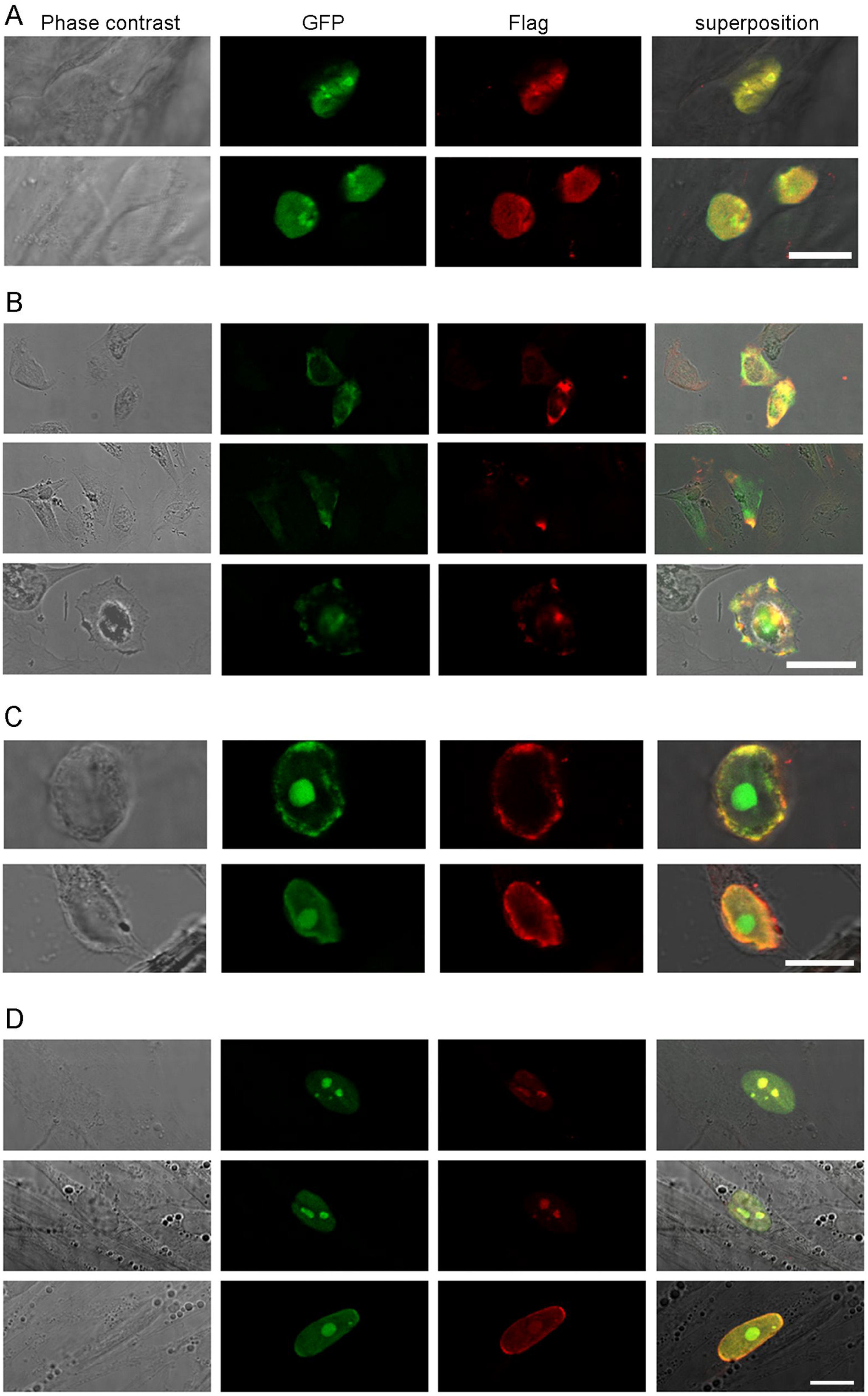

## Author Contributions

Dmitri Panteleev made experimental part such as Western Blot, 2D Western Blot, Immunofluorescent staining of cells, Rapid Amplification of cDNA Ends, Southern blot hybridization of products of 3’RACE, Dzhirgala Shamadykova cultivated cells required for research, Sergey Drozd made experimental part such as RT-PCR, Nikolai Pustogarov made experimental part such as Southern blot hybridization, Alexander Revishchin made experimental part such as Immunohistochemical analysis, Sergey Goryanov – surgical removal of the tumor and Alexander Potapov carried out management of the medical field of study. Leadership of the study: Galina Pavlova,.

## Acknowledgements

The reported study was funded by RFBR according to the research project №17-00-00157, №17-00-00158 (17-00-00162 (K)) and 18-015-00279.

